# Functional covariance modes reveal aligned fetal and neonatal brain functional connectomes

**DOI:** 10.1101/2025.07.20.665719

**Authors:** Vyacheslav R. Karolis, Jonathan O’Muircheartaigh, Grainne M. McAlonan, Tomoki Arichi

## Abstract

Spatially distributed functional networks are a fundamental property of brain organisation. While these networks are already present at full-term birth, establishing whether they exist before birth remains problematic, given the challenges inherent to in utero fMRI. Here, we introduce a seed-based functional covariance modes (FCMs) approach, which leverages inter-subject variability in connectivity between distinct regions and the rest of the brain to infer whole-brain network configurations. Unlike standard group-level independent component analysis, which consistently fails to reveal spatially distributed neural networks in fetal populations, FCMs successfully delineated a range of brain-wide functional networks in utero with high spatial fidelity to neonatal network maps, including bilateral organisation - a landmark feature of many neonatal functional networks. Furthermore, we demonstrated that bilateral fetal networks preferentially clustered along the brain midline and around cortical limbic territories. In contrast, single-hemisphere dominant networks mapped onto areas associated with hemispheric functional asymmetry in the mature brain, including the highly lateralised language circuits. Together, these findings provide coherent evidence that the blueprint for the functional brain architecture, including its functional specialisation, is formed before birth.

## INTRODUCTION

Evidence suggest that that brain functional networks first emerge very early in human development ^1,2^. At birth, functional network organisation, observed using fMRI, reveals a tendency for networks to show a near-symmetrical organisation across the hemispheres or to involve a node in the contralateral hemisphere, even in networks with a predominantly unilateral spatial layout ^3–5^. These findings are corroborated by histological^6,7^ and structural connectivity studies^8,9^ demonstrating the arrival at the cortical plate of collateral afferent thalamic inputs and callosal fibres, linking homologous regions across hemispheres, by the third trimester *in utero*.

A growing body of evidence, including those obtained in preterm born infants^1^, has begun to suggest that functional connectivity features characteristic of the neonatal brain may also be present before birth^10,11^. However, reliably demonstrating this using *in utero* fMRI is a significant challenge due to the difficulties associated with performing robust imaging studies with small, moving, and rapidly developing fetal brains. Despite advances in fetal fMRI image processing and analyses^12–15^, methods like Independent Component Analysis (ICA), a widely used and powerful tool for extracting spatial maps of resting-state networks^16,17^, have yet to demonstrate robust inter-regional connectivity features in the *in utero* brain^15,18^. Consequently, it is unclear whether long-distance - particularly, interhemispheric - connections is a feature of the brain function that emerges over the birth transition, or whether the difference between fetal and neonatal functional network organisation is due to the inherent practical and methodological differences arising from imaging a fetus in the womb versus a sleeping neonate.

The ability to robustly test if connectivity patterns are partially replicable across the birth divide using consistent methodology would thus be a critical step towards contextualisation of the fetal fMRI observations, enabling their interpretation relative to better-characterised postnatal networks. Importantly, methodological frameworks in this domain should ideally be tailored to handle the unique analytical demands of the fetal brain - specifically its rapid, almost daily, physiological transformations. In the past, we proposed an analytical framework that explicitly takes advantage of the developmental nature of the data, by leveraging cross-sectional age-related differences to infer maturational functional network properties in the fetal brain^10^. Extending this approach, here we introduce a simple and computationally inexpensive framework that leverages *full-spectrum* inter-individual variability in functional connectivity to infer whole-brain network configurations. Given its design, this approach (Fig. 1) is called *seed-based functional covariance modes* (FCMs) and belongs to a broader class of models, known as “functional covariance networks”^19,20^. Using the openly available large resource of perinatal fMRI data from the developing Human Connectome Project (dHCP)^3,15^ (see Data Availability Statement), we first test the FCMs method using the neonatal dataset to evaluate its performance against the canonical networks defined by standard group-ICA factorisation. With this in place, we turn to the fetal dataset to demonstrate that FCM can delineate a range of spatially distributed functional associations in the *in utero* brain, which align with the previously identified neonatal networks and known system-level principles of functional organisation before birth.

## RESULTS

### Neonatal dataset

The derivation of FCMs comprises 4 steps: 1) computing seed-to-brain correlation maps in native fMRI space, 2) projecting maps into a template space; 3) concatenation of individual maps into a 4D volume (aka “subject-series”^19^) and 4) independent component factorisation to obtain spatial maps representing functional networks. We start with assessment of FCMs performance in neonates, where standard (group-ICA) analyses are typically able to delineate a range of spatially distributed networks that could be used to benchmark FCMs performance. Here we used a subset of the open-source neonatal dHCP dataset (see Data Availability Statement), consisting of 311 neonates of nearly-symmetrically distributed age around the sample mean of 37.5 weeks postmenstrual age, approximately matching the age of the oldest subjects in the fetal sample^10^.

Naturally, two key parameters directly influence network factorisation: (a) dimensionality (the target component count), which dictates the trade-off between discovering novel networks and fracturing existing ones into sub-components^21^, and (b) seed location, which governs whether spatial map configurations are uniquely determined by seed placement or remain insensitive to it. We start with assessment of the FCMs computed using left thalamus as a seed and exploring two choices of dimensionality (*d*) of factorisation, *d* = 25 and *d* = 40. The full set of maps is shown in Supplementary Fig. 1 and 2; these and all other maps reported below are also available on the online resource associated with the paper (see *Data Availability Statement*). To aid assessment of the maps’ spatial layout, in particular the location of their centres-of-mass/highest values, visualisation of every spatial map in this report is scaled between *z* > 3 and its maximum value.

The results of group-ICA factorisation with *d* = 25, reported in^10^, were used for initial comparisons. In Fig. 2A these group-ICA (gica) maps are shown with the best visual matching *d* = 25 and *d* = 40 FCMs. The two approaches show high visual agreement of the identified spatial patterns, demonstrating that seed-based FCM provides a robust representation for the neonatal group-level networks if gica maps are taken as a benchmark. These not only include patterns which are good candidates for representing assumed neurally determined connectivity patterns but also patterns likely to be driven by other physiological effects including vascular signal (cf. patterns paired with gica 17, 22, 24).

Several qualitative features appear to stand out following a visual inspection. Firstly, FCMs factorisation both tended to be more sensitive to noise, manifested by isolated, non-clustered voxels scattered across the brain while capturing the core network nodes, and also created a larger number of components (approximately, 50%) that could be unambiguously classified as structured noise of a vascular origin. Secondly, both *d* = 25 and *d* = 40 FCMs had a greater propensity for bilateral patterns, i.e., spatial maps that were near-symmetrically distributed between left and right hemispheres, specifically in occipital (visual) and peri-insular (auditory) cortices. Finally, the shift in FCMs dimensionality from 25 to 40 components did not result in losing distributed spatial characteristics observed in the group-ICA maps, nor the ratio between components of apparent vascular vs. neural origin changed, suggesting that d = 25 FCMs may provide insufficient description for the richness of functional networks in the neonatal brain.

Next, in several analyses, we assessed the importance of the seed site location. Firstly, we found that using the right thalamus as a seed rendered comparable results to those for left thalamus (Supplementary Fig. 3 and 4). As left and right thalami can be closely related in the brain’s functional organisation, and consequently, reveal similarities in their correlation maps, we further repeated the analyses using seeds in the left cerebellum. Cerebellum-based FCMs (Supplementary Fig. 5 and 6) tended to recover either the same spatial pattern (with differences in fine details of their spatial layout) as the thalamus-based FCMs or differing by their unilateral vs. bilateral interpretation. Thus, the choice of a specific seed had implications for the specificity of the derived spatial maps, not their global configuration. This indicates that combining several seeds in the analyses may boost SNR by reinforcing repeating patterns in the correlational structure and therefore efficiently deal with its increased sensitivity to noise compared to group-ICA.

To assess this possibility, we computed multiregional FCMs (mrFCMs), using the left and right thalami and cerebellum together as seeds. Maps for the left part of each anatomical structure were left-right mirrored and the averages between the right and mirrored left maps were computed. (Note that these maps become left-right equivalent). Tensor ICA concatenation, as implemented in melodic FSL^22^, was used to combine the left-right average maps for thalamus and cerebellum in a unified analysis. The effect of each type of the data fusion (left-right averaging vs tensor-ICA concatenation) on the results were assessed later in detail in the context of the fetal dataset, where the gains from boosting SNR can potentially be more evident. The selected maps for *d* = 40 mrFCMs, matching the group-ICA networks, are shown in Fig. 1B and their complete set is shown in Supplementary Fig. 7. As expected, compared to single-seed FCMs maps, mrFCMs appeared to provide better noise suppression while preserving the spatial specificity of single-seed FCMs and sometimes providing better visual matching to gica maps than single-seed FCM (cf. maps paired with gica maps 2,3,12, 15 in Fig. 1A).

**Fig. 1.**
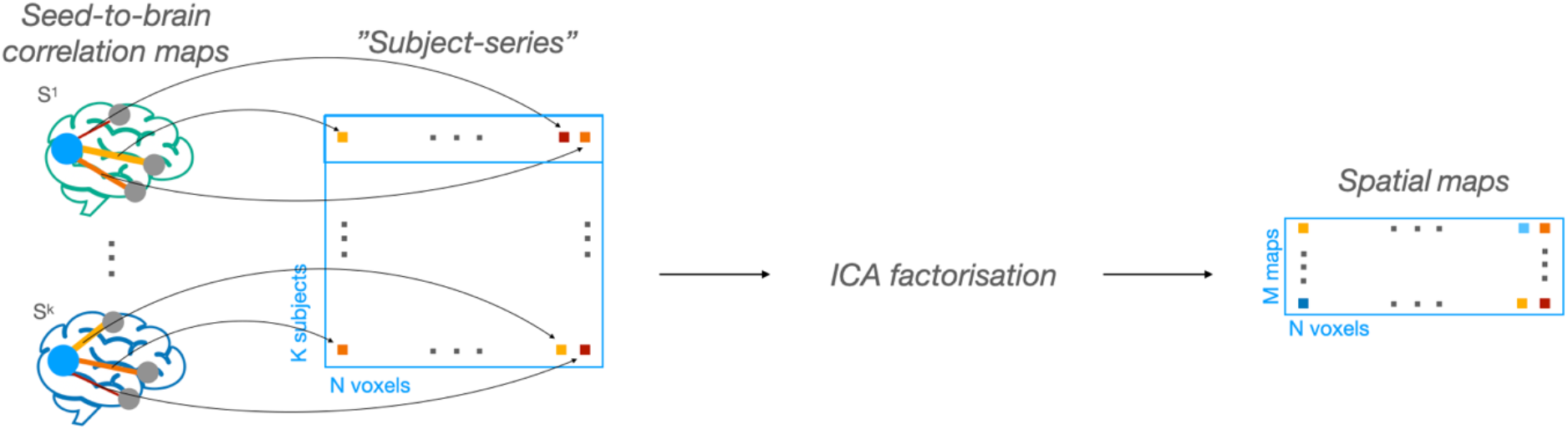
Derivation of seed-based functional covariance modes

**Fig. 2.**
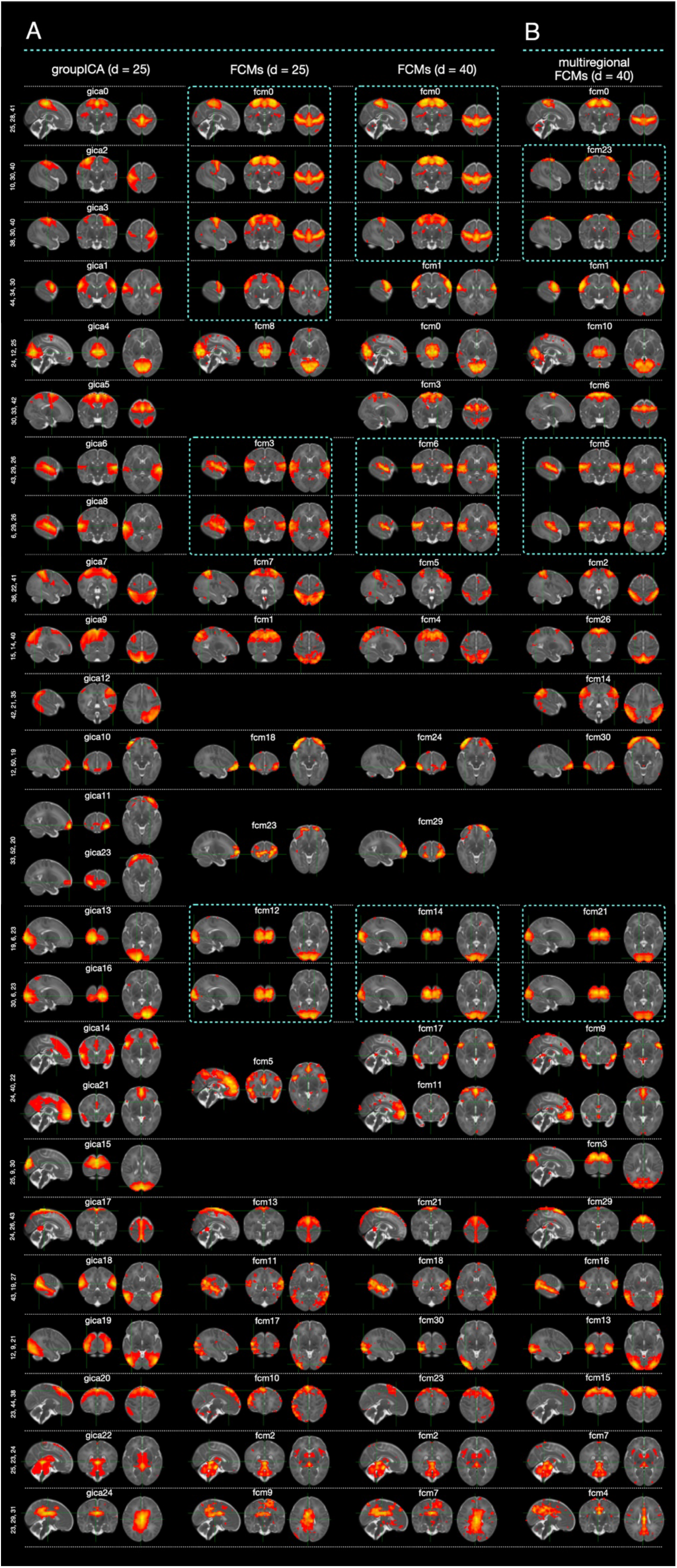
Similarity between FCMs and d = 25 group-ICA results. A) group-ICA and FCMs seeded with left thalamus. Images are shown in radiological orientation (Left is right). B) multiregional FCMs. Images are left-right ambiguous

The final part of the analysis in the neonatal dataset aimed to assess whether FCM factorisation aligns with group ICA factorisation as the dimensionality of both models increases. In other words, we explored if the observed differences between d=25 gica and FCMs maps are not due to the patterns “invented” by the FCM approach but rather are in line with the variations that can occur within the group-ICA approach when the latter is performed using a different choice of dimensionality. For this, the group-ICA factorisation was obtained using *d* = 40 (Supplementary Fig. 8). The results indicate a stable alignment between the two approaches: among 5 non-trivial mrFCM patterns (i.e., those which spatial layout suggest its biological veracity), that could not be matched to any of *d* = 25 gica maps, 4 could be reliably paired with *d* = 40 gica maps (Fig. 3).

**Fig. 3.**
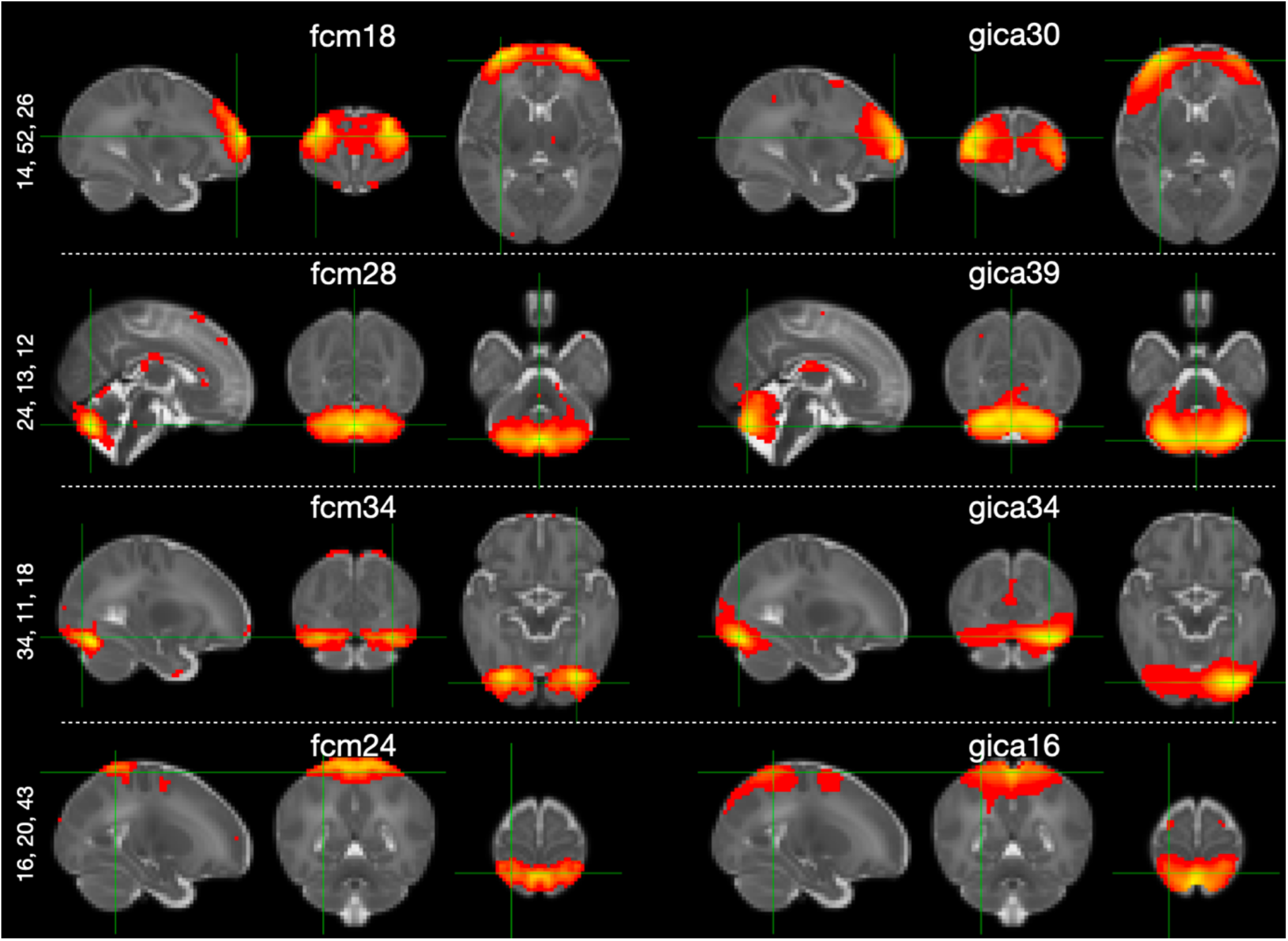
Effect of increased dimensionality. d = 40 mrFCMs maps that that could not be matched to any of d = 25 group-ICA but can be paired be d = 40 group-ICA results. FCMs are left-right ambiguous. Left is right for gica maps.

### Fetal dataset

Using the neonatal dataset, we demonstrated that the seed-based FCM approach can produce a valid spatial factorisation when group-ICA results are used as a benchmark. We now turn to the analyses of the fetal data, which have proved to be very challenging for recovery of spatially distributed functional brain networks using conventional methods.

The analyses were run on the sample of 201 fetuses from the dHCP cohort, aged 24.5 – 38.29^15^. We started by considering the configurations of the *d* = 40 multiregional FCMs. The results are shown in Fig. 4, where they are graphically organised to enable an easier identification of their anatomical location. These results demonstrated that mrFCMs represent a more powerful tool for extracting spatially distributed patterns of functional connections than the conventional group-ICA approach (see Supplementary Fig. 9 for d=40 fetal group-ICA). Notably, FCMs displaying symmetrical organisation or secondary, non-negligible nodes in the contralateral hemisphere were particularly prevalent. Additional analyses demonstrated that the left-right mirror-symmetric re-enforcement represents a critical factor for the ability to derive spatially-distributed patterns in the fetal data, as shown by comparison of left+right averaged thalamus-based FCMs (Supplementary Fig. 10) and FCMs obtained by combining left thalamus and cerebellum seeds (Supplementary Fig. 11).

**Fig. 4.**
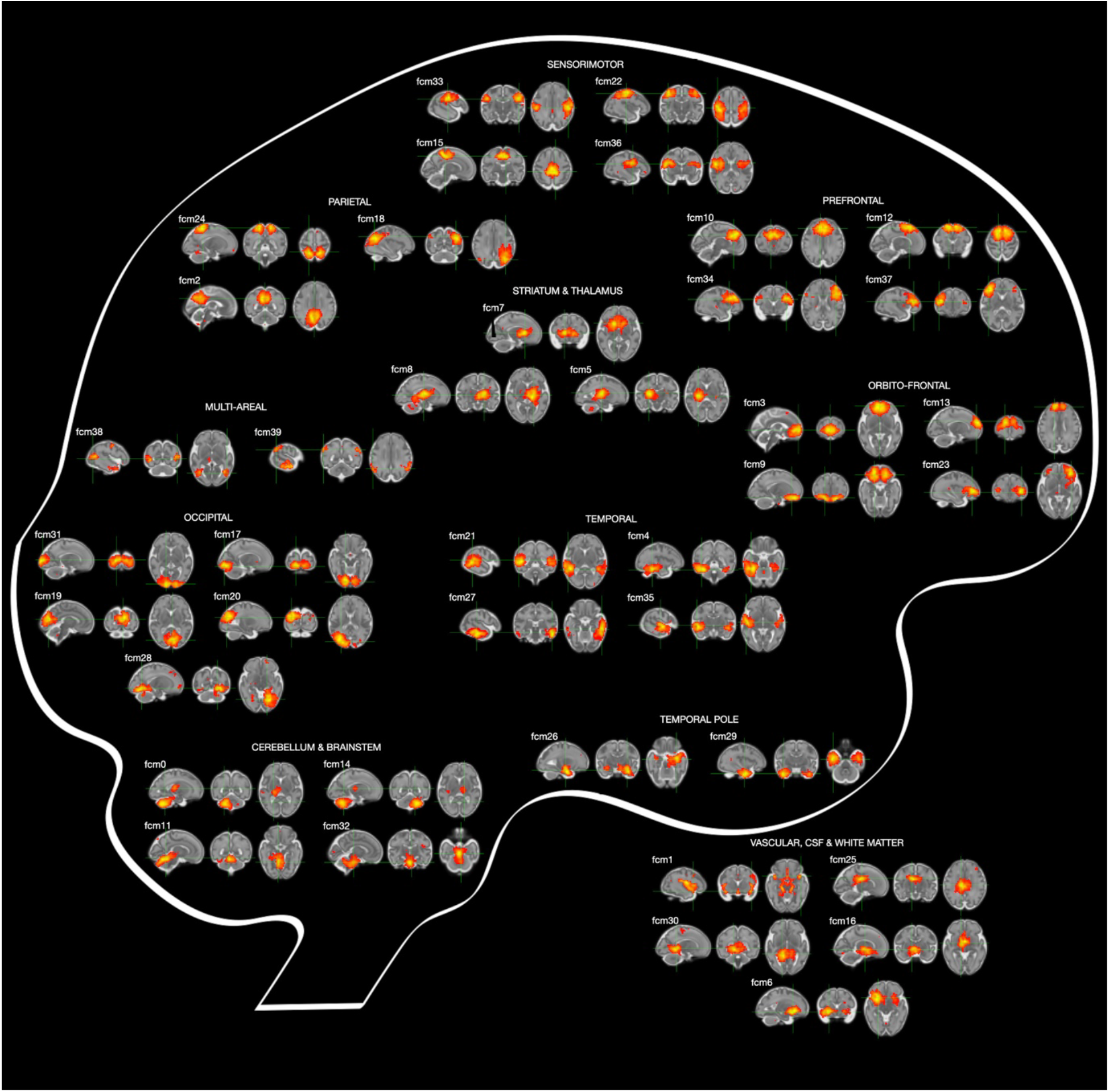
d = 40 fetal mrFCMs. Images are left-right ambiguous.

We next compare the fetal mrFCMs with neonatal *d* = 40 group-ICA networks. Fig. 5A shows the mrFCMs which either very accurately match the *d* = 40 neonatal gica maps or represent their *multi-nodal* sub- or super-part. Beyond that, we also observed that a global vascular-related component (fcm1) included areas which overlapped with isolated neonatal gica maps of a likely neuronal signal, namely peri-insular/auditory gica 5 and gica 7 and gica 31 (salience network). Other correspondences include two unilateral cerebellar components (cf. bilateral neonatal gica 39 in Fig. 3) and additional matching patterns between neonatal group-ICA and d =25 fetal mrFCMs factorisation (Supplementary Fig. 12; e.g., fcm 20 vs. gica 34, fcm16 vs. gica 5 and 7, fcm 9 vs. gica 17). A closer inspection of the maps that were undeniably matching between neonates and fetuses is instructive to appreciate the differences in the signal characteristics, with the centres-of mass appearing to be biased towards veins in the neonatal maps and towards white matter in the fetal maps (Fig. 5B).

**Figure 5.**
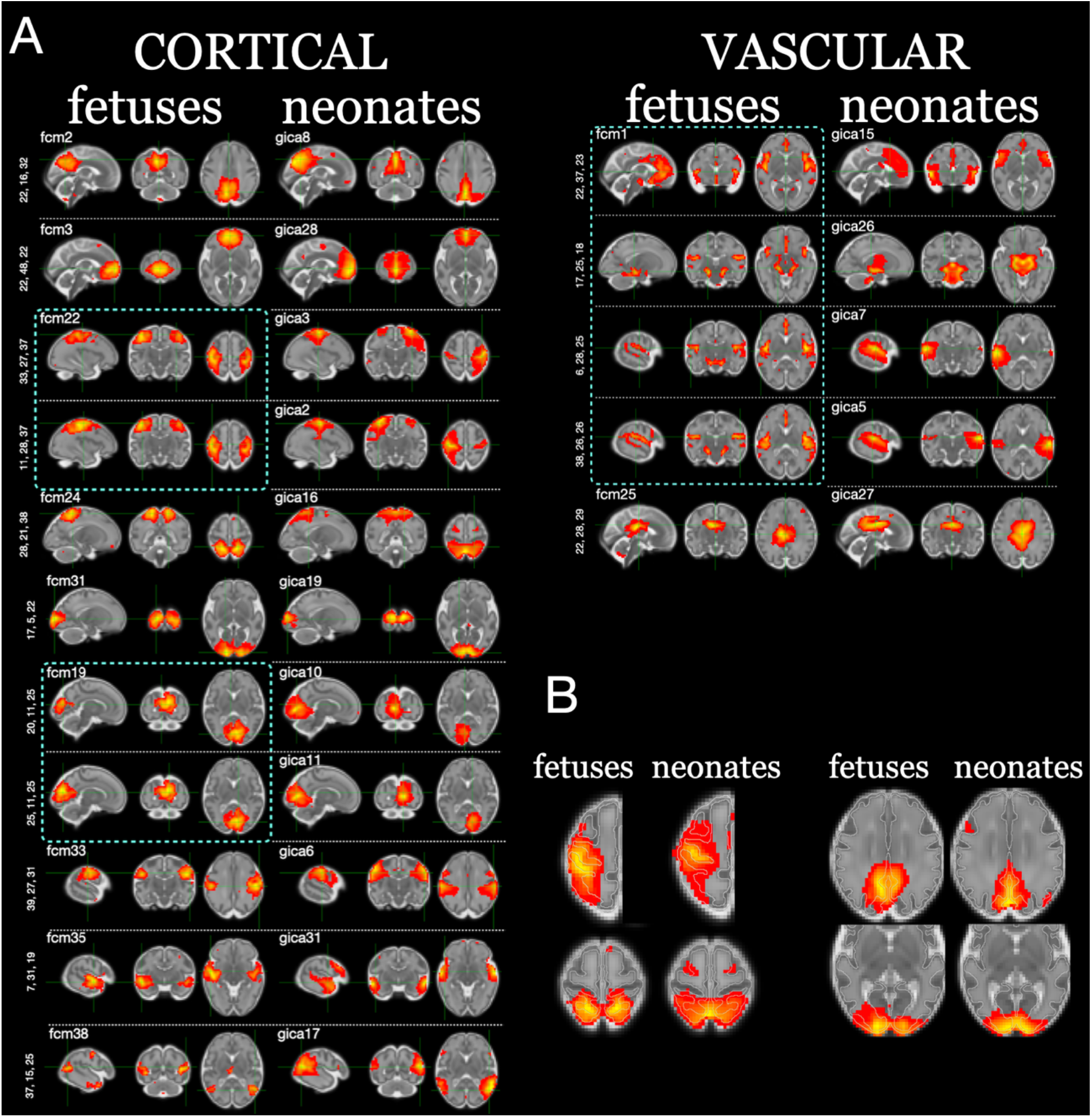
Comparisons between d = 40 fetal FCMs and neonatal d = 40 group-ICA maps. A. Matching patterns; B) Signal characteristics in fetal and neonatal maps.

The analysis of neonatal dataset indicated that an increase from *d* = 25 to *d* = 40 did not have a detrimental effect on the ability to derive spatially distributed patterns but improved their spatial specificity; consequently, an even higher dimensionality may potentially permit a more efficient decoupling of such patterns from a background covariance structure dominated by local smoothness in the correlational maps. This proved to be the case, as high dimensional (d = 80) factorisation revealed a variety of collateral patterns (Data availability, Resource 1). In Fig. 6 we combined the maps that showed a left-right near-symmetric organisation vs the maps that showed single-hemisphere dominance. The FCMs with bilateral patterns preferentially mapped along the midline separating the two hemispheres, both dorso-laterally and medially. The exceptions to this pattern were cortical limbic areas, combining ventral orbito-frontal and temporal pole regions; ventral primary sensorimotor cortices; regions along the mid-to-posterior middle temporal sulcus. In contrast, single-hemisphere dominant regions preferentially mapped onto lateral cortical territories - that included inferior prefrontal cortex, posterior superior temporal gyrus, auditory cortex, middle pre-frontal cortex, angular gyrus, lateral sensorimotor regions, frontal and parietal eye fields, lateral anterior temporal and lateral occipital cortices – and medial ventral regions (fusiform, lingual and parahippocampal gyri).

**Fig. 6.**
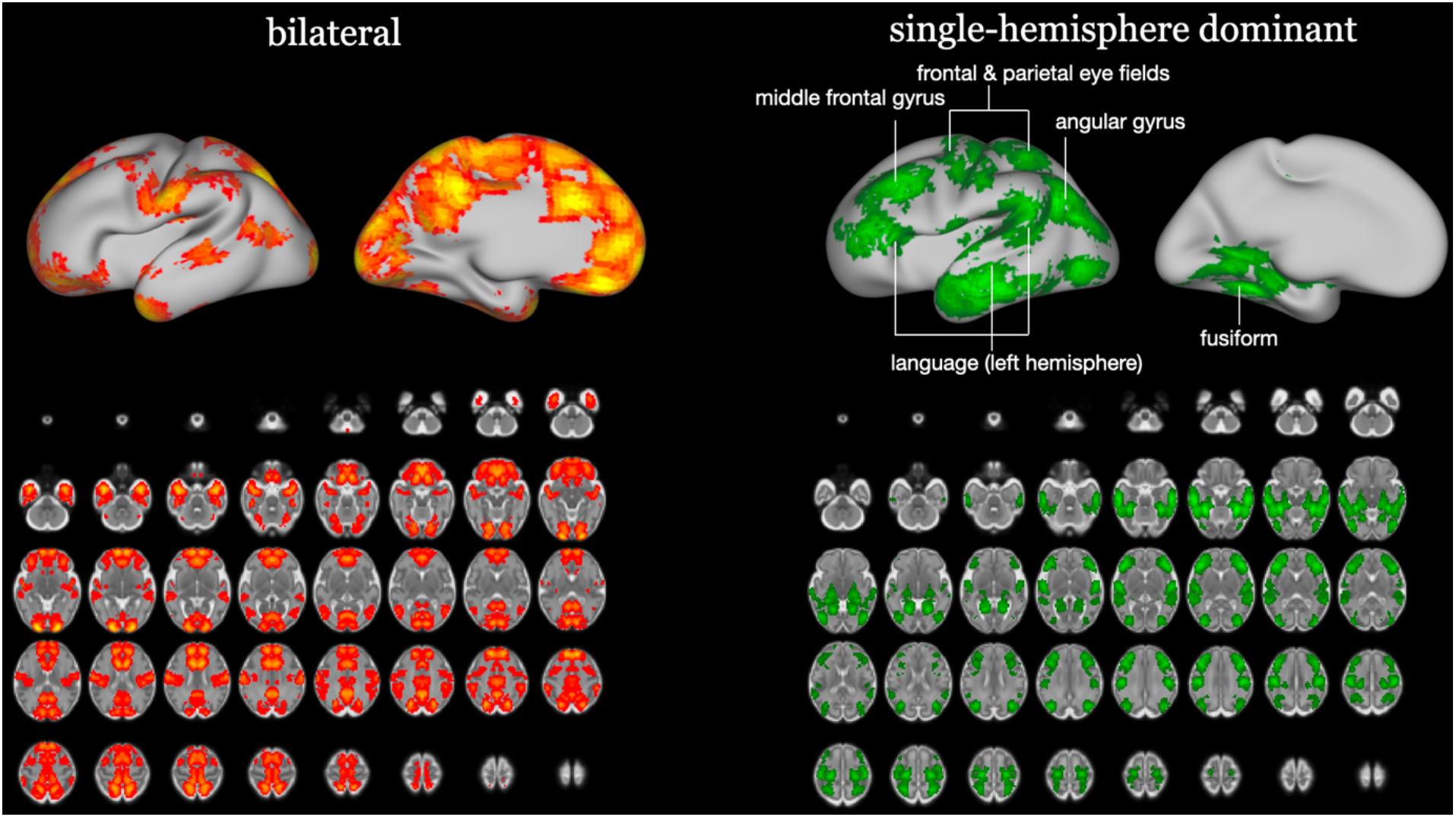
Spatial distribution of bilateral and single-hemisphere dominant maps in d = 80 FCMs. Images are left-right equivalent and were produced by combining, separately, near-symmetric and predominantly unilateral z-scored spatial maps, followed by their left-right flipping and extracting the maximum value across all maps and both their flipped and non-flipped versions. The resulting maps were thresholded at z = 5 and shown both in the volumetric space and the cortical surface, to aid visualisation of the distribution of bilateral and unilateral patterns.

## DISCUSSION

In this paper, we introduce an analytical technique that utilises inter-individual differences in seed-based connectivity to infer the brain’s functional network characteristics. The motivation for developing this approach builds on our earlier work^10^ showing the utility of inter-subject variability for the inference of network architecture in the fetal brain, and is driven by the primary objective of defining network-like properties in this population. Based on its underlying structure, we term this approach seed-based functional covariance modes (FCMs) and attribute it to a broader and relatively overlooked class of models known as functional covariance networks^19,20^.

We first assessed FCMs performance in the context of neonatal fMRI data. We demonstrated:1) factorisation of network-like properties comparable to factorisation using the conventional and widely-used group-ICA approach, and 2) relative tolerance to the choice of the seed location, which allows combining maps derived using different seeds to reinforce repeating patterns in the correlational structure. Based on these results, we developed an optimised strategy, *multiregional FCMs*, to approach the fetal dataset.

We showed that - in the context of fetal fMRI - FCMs are not just an inconsequential alternative to the standard group-ICA maps; rather they mark fetal fMRI as a key application for functional covariance models to detect network-like functional brain properties with higher efficiency. Importantly, for the first time and despite the fundamental differences in the signal characteristics between fetal and neonatal imaging of function, we were able to detect spatially distributed network-like patterns which could unambiguously be matched to the patterns observed in the neonatal brain using group-ICA, thereby boosting confidence in the biological veracity of observations made using fetal fMRI. A substantial proportion of these patterns localised to primary and secondary cortical areas, supporting current models in which these regions anchor the earliest stages of functional maturation^23,24^. Their demonstration has been a lingering problem for the fetal fMRI field^10,15^, as the ambiguous spatial characteristics of the fetal functional network maps, obtained using other methods, undermined their biological credibility, complicate their interpretation, and hinder generation of hypotheses about fetal functional brain development.

Furhtermore, FSMs identified a broad range of bilateral patterns in the fetal brain, which crucially recapitulate^25^ the dominance of interhemispheric relationships characteristic of neonatal networks^4,5^. These patterns were particularly prominent with high-dimensional factorisation, which was an unexpected finding, given that increase in the dimensionality of factorisation in the standard (group-ICA) settings oftentimes results in the fractionation of distributed patterns into sub-parts. The most prominent motif was the higher prevalence of the bilateral patterns along the brain midline, comprising the medial and dorsal regions of the brain, and consistent with the organisation of callosal connectivity in the human brain^26^ as well as with the previously reported fMRI results obtained in preterm babies^1^. There were prominent exceptions to this global structure, in particular encompassing the mouth area of sensorimotor area, posterior regions of temporal cortex, which in adult brain are associated with social interaction, and limbic ventral orbito-frontal cortices/temporal pole regions. This potentially points to a developmental hierarchy prioritising the brain systems needed to ensure survival at early stages of life^27^. In contrast, among regions which showed single-hemisphere dominance we observed the areas which, in the adult brain, are associated with heavily lateralised (in the left hemisphere) language function. Functionally related anterior lateral temporal cortex, involved in the semantic processing^28,29^, was also present, as well as fusiform, parietal and frontal eye field, parahippocampal and middle portion of the motor cortices, all known to demonstrate a high degree of hemispheric functional specialisation^26^. The middle frontal gyrus, another region present in this single-hemisphere dominant map, was also hypothesised to play a distinctive functional role across hemispheres: the connector of the two major attentional systems^30^, top-down dorsal and bottom-up ventral^31^, in the right hemisphere and closely linked to the language networks in the left hemisphere, implementing the executive control component of the language function. Together, our findings provide coherent evidence that the blueprint for the distributed functional brain architecture, including its hemispheric functional specialisation, is formed before birth.

The utility of inter-subject variability for the analyses of fetal brain network architecture was highlighted in our previous work^10^ that leveraged age-related differences to infer network organisation in the fetal data. The method, called matnets, can therefore also be considered a special case of the functional covariance network approach. Compared to that, FCMs combine improved power in detecting spatially distributed patterns with a significantly lower computational cost (linear vs. quadratic scaling of computation times). Methodologically, our approach is related to the two approaches which originally introduced functional covariance networks to the neuroscientific community. Zhang et al.^20^ introduced seed-based functional covariance networks, where a particular seed was used to reconstruct a network it belongs to. Taylor et al.^19^ then used a combination of within-voxel signal metrics (e.g., temporal deviations or amplitudes) and ICA factorisation to reconstruct networks that proved to be similar to the results of the standard group-ICA. Our approach lies at the intersection of these two approaches: in contrast to Taylor et al. our approach is grounded in the correlational structure, i.e., relying on the statistical contrast that underpins the mainstream network analysis methods; in contrast to Zhang et al., we show that the use of a particular seed does not prevent whole brain network organisation inference, at least in the perinatal cohorts tested here.

The alignment of network patterns between cohorts is particularly striking given the profound differences in underlying signal characteristics, which extend well beyond disparities in signal-to-noise ratio. Thus, we note that the neonatal maps have a tendency to be organised along sulcal grooves, a likely result of the venous bias of gradient-echo imaging^32,33^. This factor is also likely to contribute to overly smooth appearance of the neonatal networks, potentially leading to overestimation of their spatial spread. In contrast, the fetal functional network maps (both group-ICA and FCMs) demonstrate a spatial bias towards infra-cortical white matter. This factor has previously been noted by us in relation to the dHCP dataset^10^ and also appears to manifest itself in results derived using other datasets^11,34^. Here, this bias may further be exacerbated by a longer echo times, utilised to provide more efficient suppression of the signal emanating from fatty maternal tissues and, consequently, the dominance of white matter contrast in the images due to its longer T2* (this can be ascertained by comparing grey-white matter contrast in fetal and neonatal dHCP datasets^3,15^). The different spatial biases are likely to cause a shift in network estimated locations and consequently, may conceal further potential similarities between fetal and neonatal network organisation.

## Limitations

### Two limitations have to be acknowledged

Firstly, a demonstration of multi-regional associations in the fetal brain required deployment of signal reinforcing strategies, the most powerful of which was left-right mirror-symmetric reinforcement. Note that this strategy reinforces mirror-symmetric patterns in any direction with respect to the seed location, i.e., it is not numerically biased to the detection of collateral patterns and therefore cannot explain their abundance due to artificially induced dependencies. The cost of such a strategy is of course the ambivalence of the spatial maps with respect to left-right orientation, that is, they are left-right equivalent. However, this does not affect the interpretation of the near-symmetric patterns, which by default are left-right ambiguous and thus small variations in their layout across the two sides of the brain can probably be ignored or averaged out. For unilateral or inter-areal patterns, given the unprecedented challenges associated with imaging brain function in utero, the importance of detecting these patterns in the context of fetal fMRI may outweigh the importance of their attribution to a specific (or either) hemisphere; the future research could potentially consider how to retrospectively map these patterns on the brain in a lateralisation-aware manner.

Secondly, a key remaining question concerns the exact nature of the information leveraged by the FCM approach to achieve a coherent network factorisation. However, unpacking the mechanism behind the FCM approach is inseparable from another question: What information is actually critical in the typical group-level analysis settings, as exemplified by group-ICA? Here, network factorisation relies on the correlation structure of fMRI timeseries within individuals, emphasising their shared, population-wide characteristics^17^. In contrast, FCM explicitly utilises inter-subject variability, clustering the brain areas based on how seed-to-brain correlation maps differ between individuals. The striking concordance observed here between FCM and group-ICA factorisations in neonates, relying on ostensibly contrasting signal characteristics, prompts a critical re-evaluation of mainstream network inference. It suggests that inter-individual variability is not merely “noise” around a group mean but is structurally aligned with the group-level architecture itself.

## METHODS

### Data

We used the pre-processed fMRI data included in the open-access release of the multimodal perinatal data by the developing Human Connectome Project Consortium^35^. The fetal dataset consists of 201 cases, aged > 24.5 gestation weeks, which data passed the quality control as described in^15^. The neonatal dataset consists of 311 cases, described in^10^ and includes all cases which gestation age is < 37 weeks and a randomly selected subsample such that the distribution around gestation age of 37 weeks has a near-symmetrical shape.

### Image preprocessing

The preprocessing stages for fetal and neonatal datasets are, respectively, described in^15^ and^3^. The warps between native and non-symmetrical template spaces are also provided by dHCP. These were concatenated with non-symmetrical to symmetrical template warp, to provide a direct mapping between native functional spaces and symmetrical template (available online, see Data availability statement).

The data from both datasets were pre-processed in an analogous way. The fMRI volumes were smoothed in the native space. The individual structural multiregional segmentations released by the dHCP (draw-EM for neonates^36^ and BOUNTI for fetuses^37^) were projected into native functional space to create a mask for a seed. The individual maps representing correlation between the mean seed BOLD signal timecourse and timecourses of each voxel in the brain were computed in the native functional space. These maps were then projected to the symmetrical group space, 40-week-old neonatal template for neonatal dataset (avialable at: https://git.fmrib.ox.ac.uk/seanf/dhcp-resources/-/blob/master/docs/dhcp-augmented-volumetric-atlas-extended.md) and 36-week-old fetal template for the fetal dataset^38^. The data across each dataset were concatenated into 4D map (“subject series”) and submitted to FSL melodic^39^ factorisation of desired dimensionality. When it was necessary to visualise together the results of fetal and neonatal analyses, the group-level neonatal maps were projected into the fetal template space.

## Data availability statement

The fMRI data reported used for the present analyses were released by dHCP, https://nda.nih.gov/edit_collection.html?id=3955. All other materials will be deposited in the following repository: https://gin.g-node.org/slavakarolis/seed-based_FCMs/main This includes:

1. All derived functional network maps reported in the paper.
2. Symmetric and fetal neonatal templates.
3. Native functional to symmetric template transformations.

## Acknowledgements

The data used in this study were acquired with the funding by European Research Council under the European Union’s Seventh Framework Programme (FP7/20072013)/ERC grant agreement no. 319456 (dHCP project). The results leading to this publication have received funding from the Innovative Medicines Initiative 2 Joint Undertaking under grant agreement No 777394 for the project AIMS-2-TRIALS. This Joint Undertaking receives support from the European Union’s Horizon 2020 research and innovation programme and EFPIA and AUTISM SPEAKS, Autistica, SFARI. JOM was funded by a Sir Henry Dale Fellowship jointly by the Wellcome Trust and the Royal Society (206675/Z/17/Z). TA was funded by a MRC Senior Clinical Fellowship [MR/Y009665/1] and receives support from the MRC Centre for Neurodevelopmental Disorders, King’s College London [MR/N026063/1]. GMM is part-funded by the National Institute for Health and Care Research (NIHR) Maudsley Biomedical Research Centre (BRC). The views expressed are those of the authors and not necessarily those of the any above-mentioned funder, the NIHR or the Department of Health and Social Care.

## Notes

### Competing Interest Statement

The authors have declared no competing interest.

### Summary of Updates

Improved flow, focus and interpretation, removed parts which are not important or redundant for the main aims.

