## Supplementary Figures for "Functional covariance modes reveal aligned fetal and neonatal brain functional connectomes"

*Figure S 1. d = 25 neonatal FCMs seeded with left thalamus.*

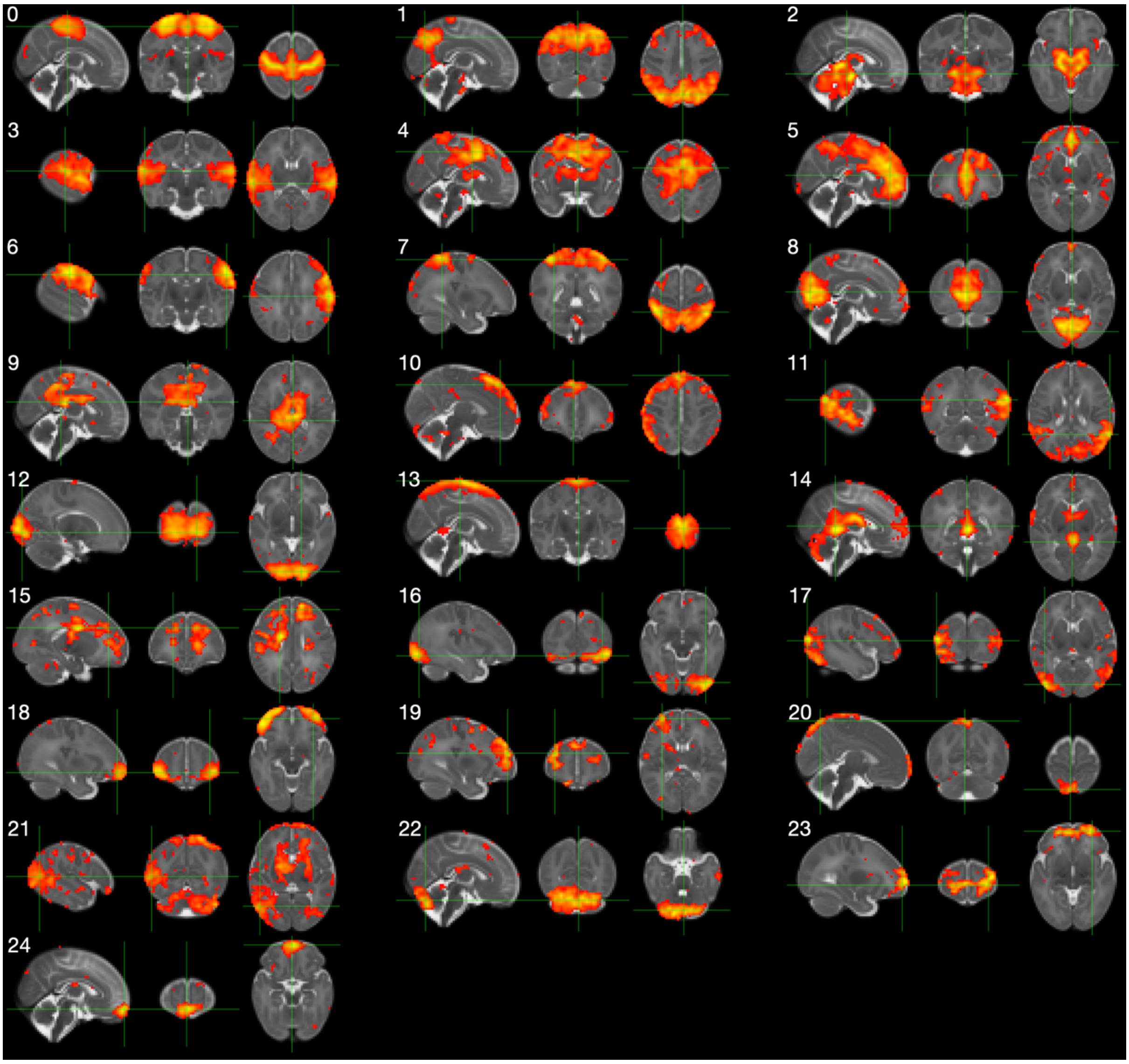

*Figure S 2. d = 40 neonatal FCMs seeded with left thalamus.*

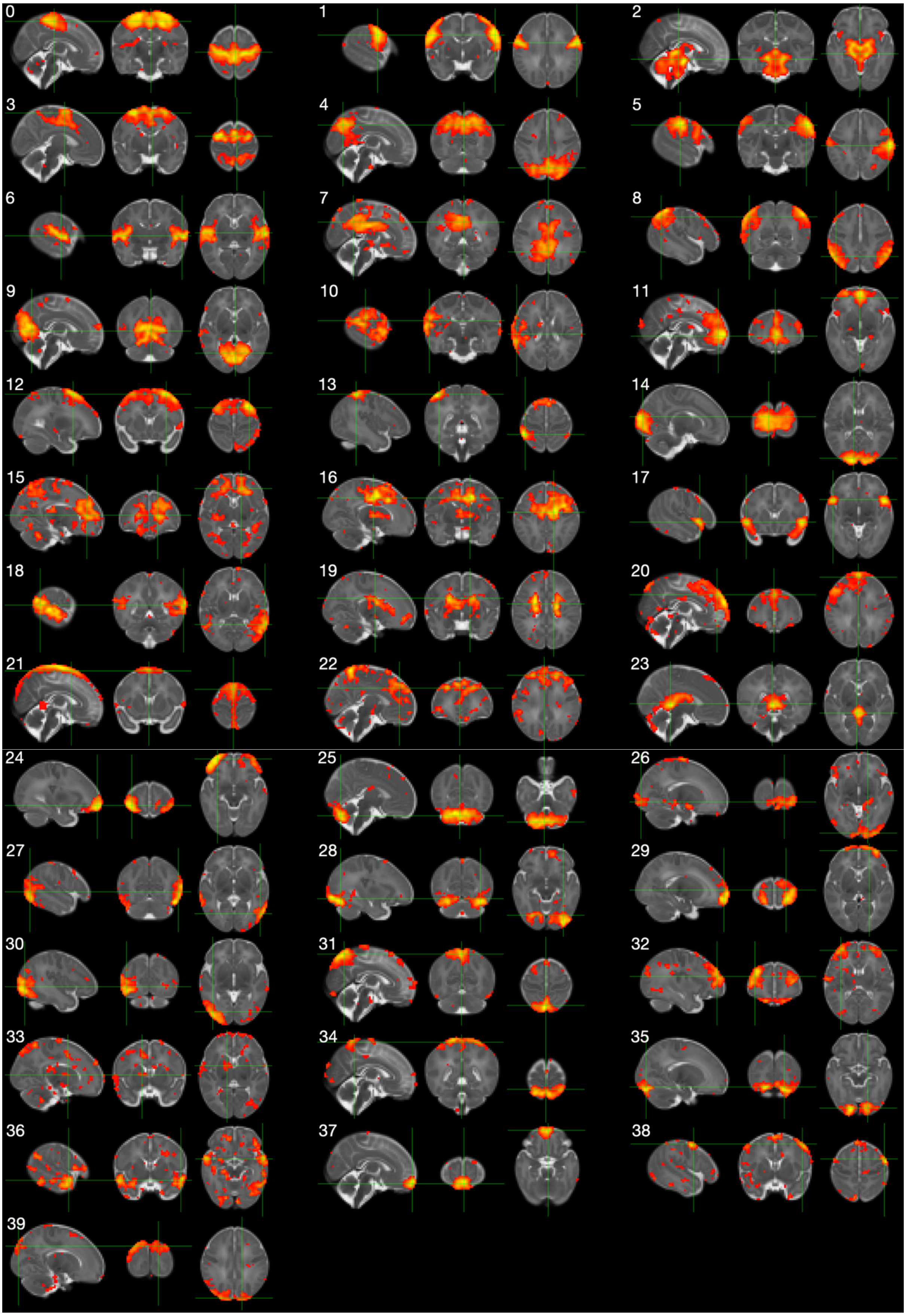

*Figure S 3. d = 25 neonatal FCMs seeded with right thalamus.*

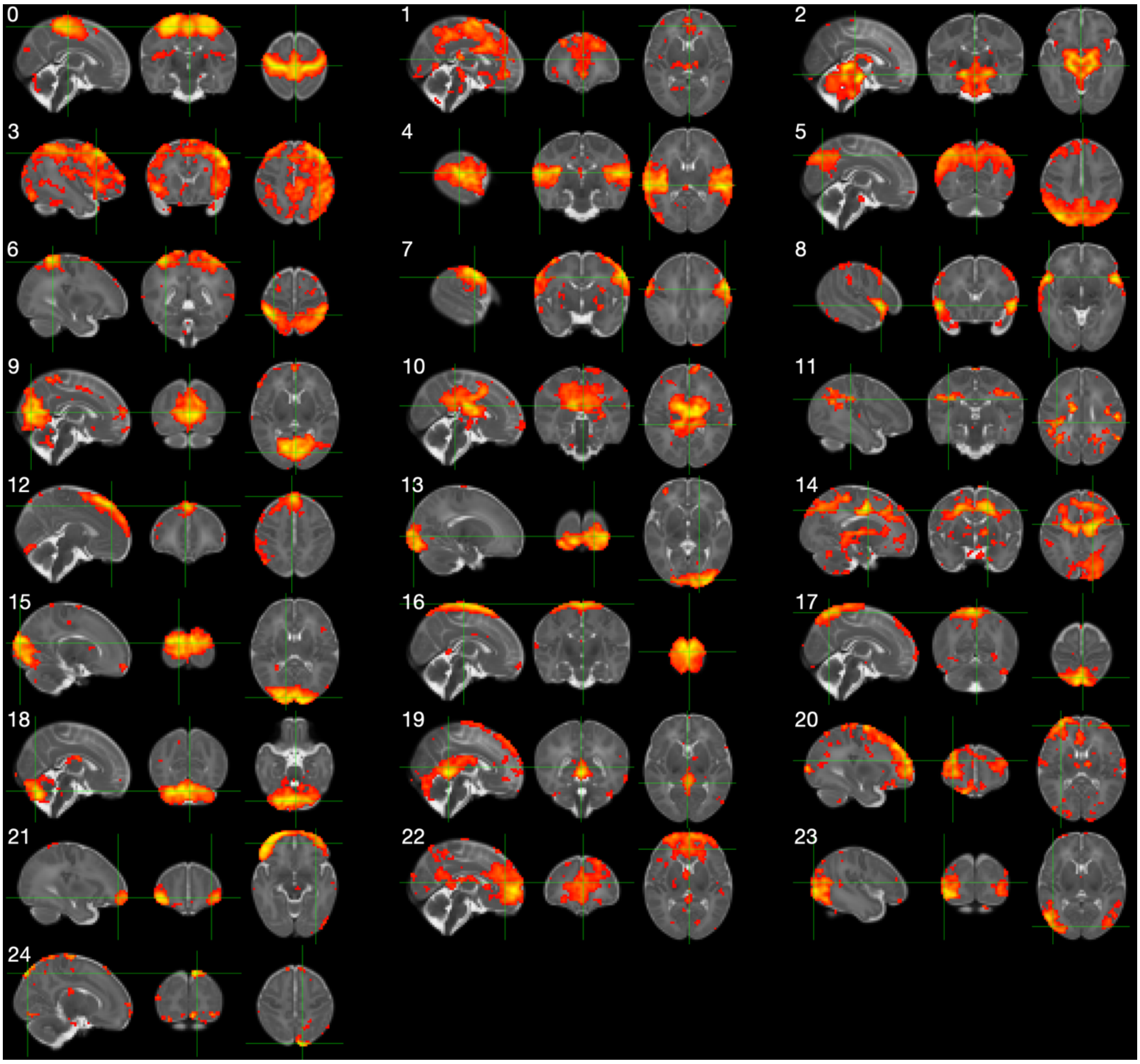

*Figure S 4. d = 40 neonatal FCMs seeded with right thalamus.*

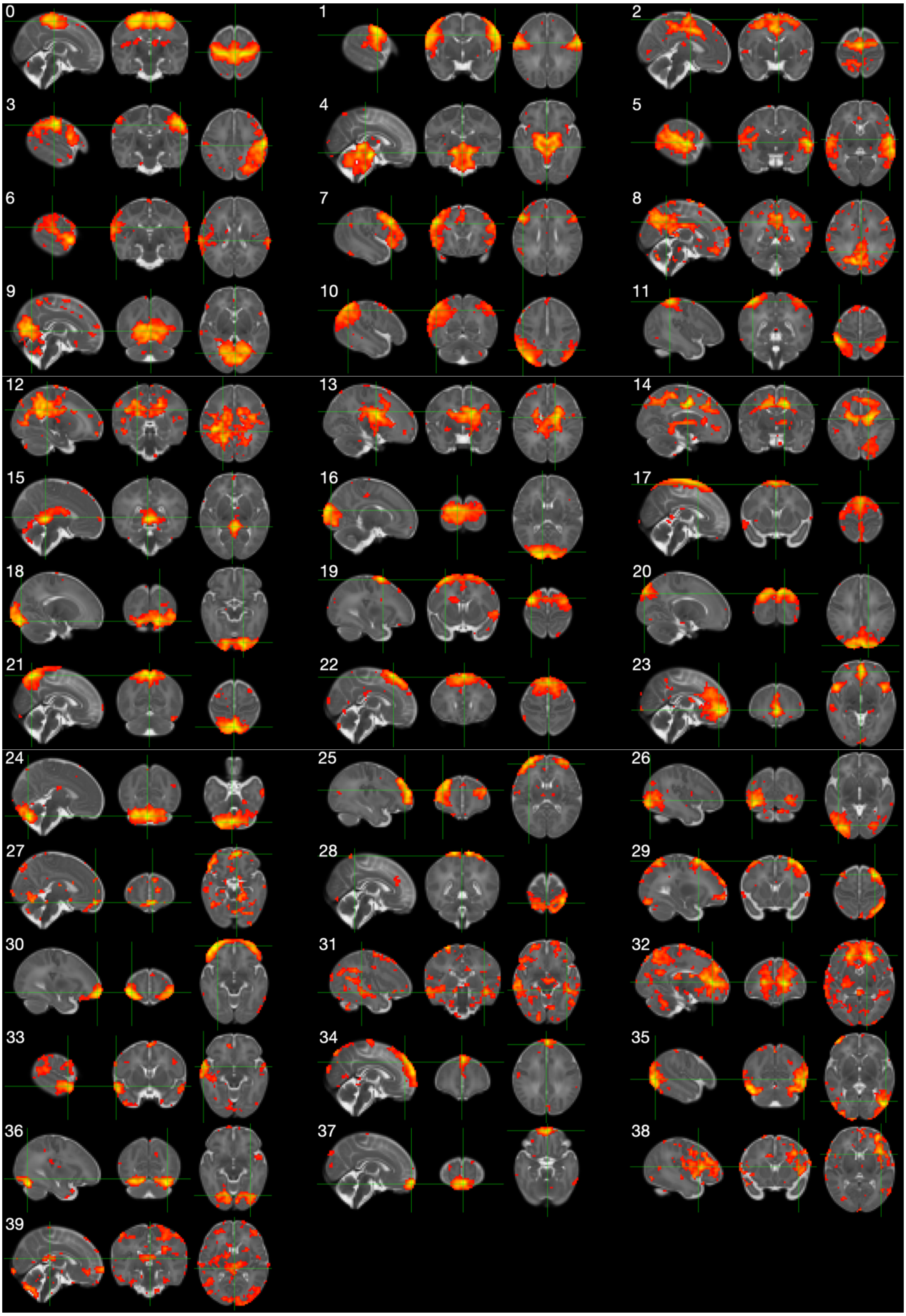

*Figure S 5. d = 25 neonatal FCMs seeded with left cerebellum.*

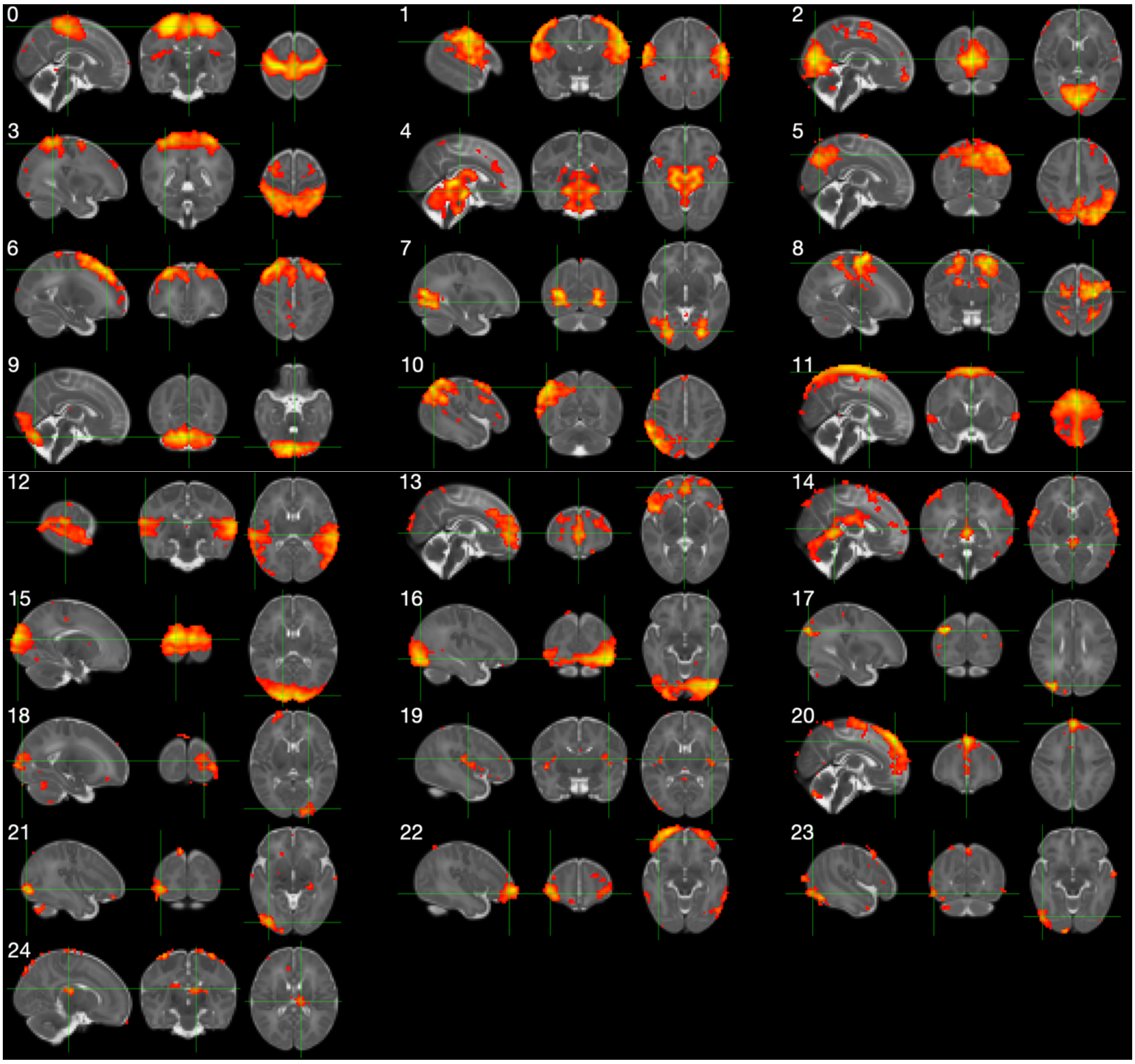

*Figure S 6. d = 40 neonatal FCMs seeded with left cerebellum.*

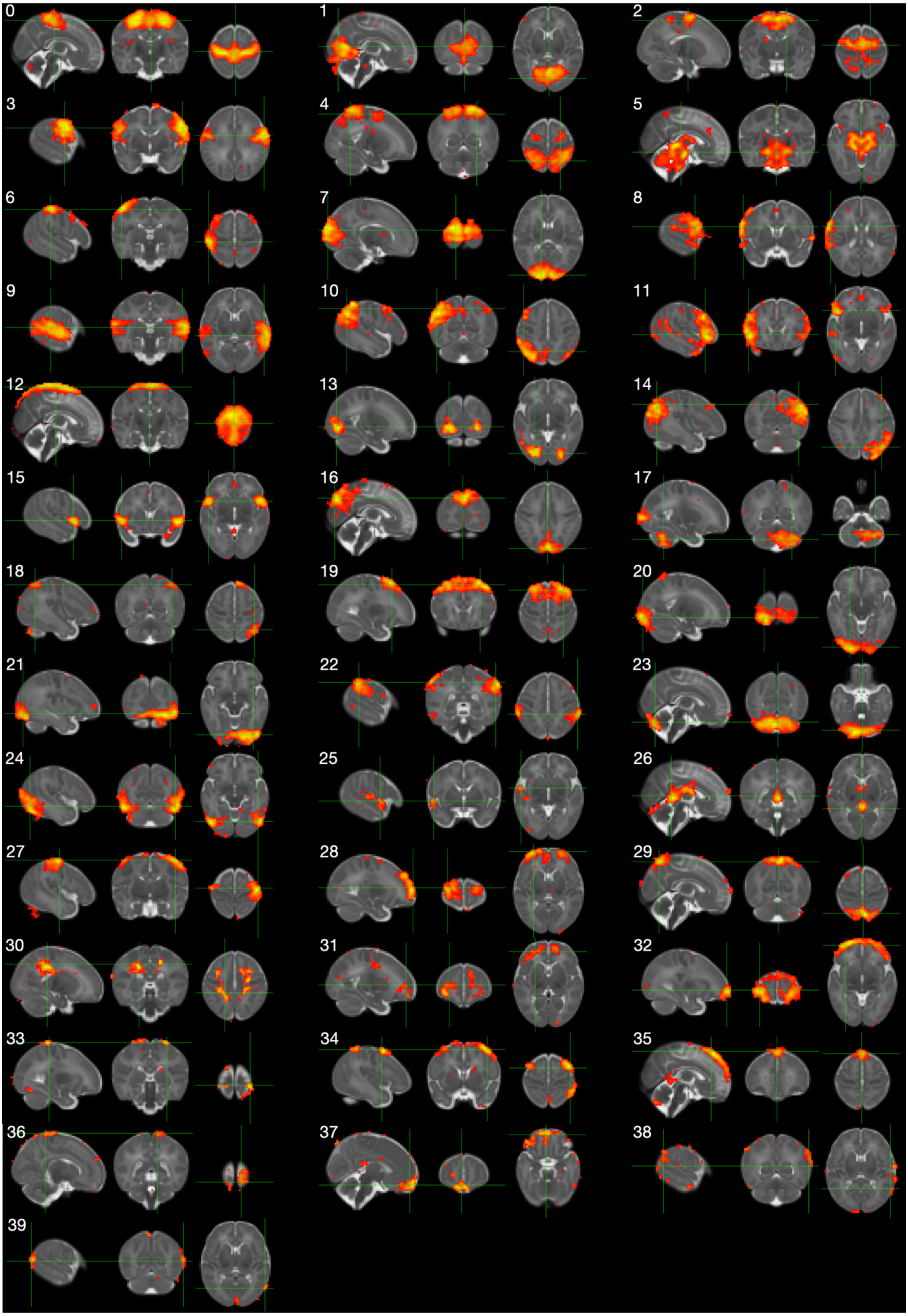

*Figure S 7. d = 40 neonatal multi-regional FCMs.*

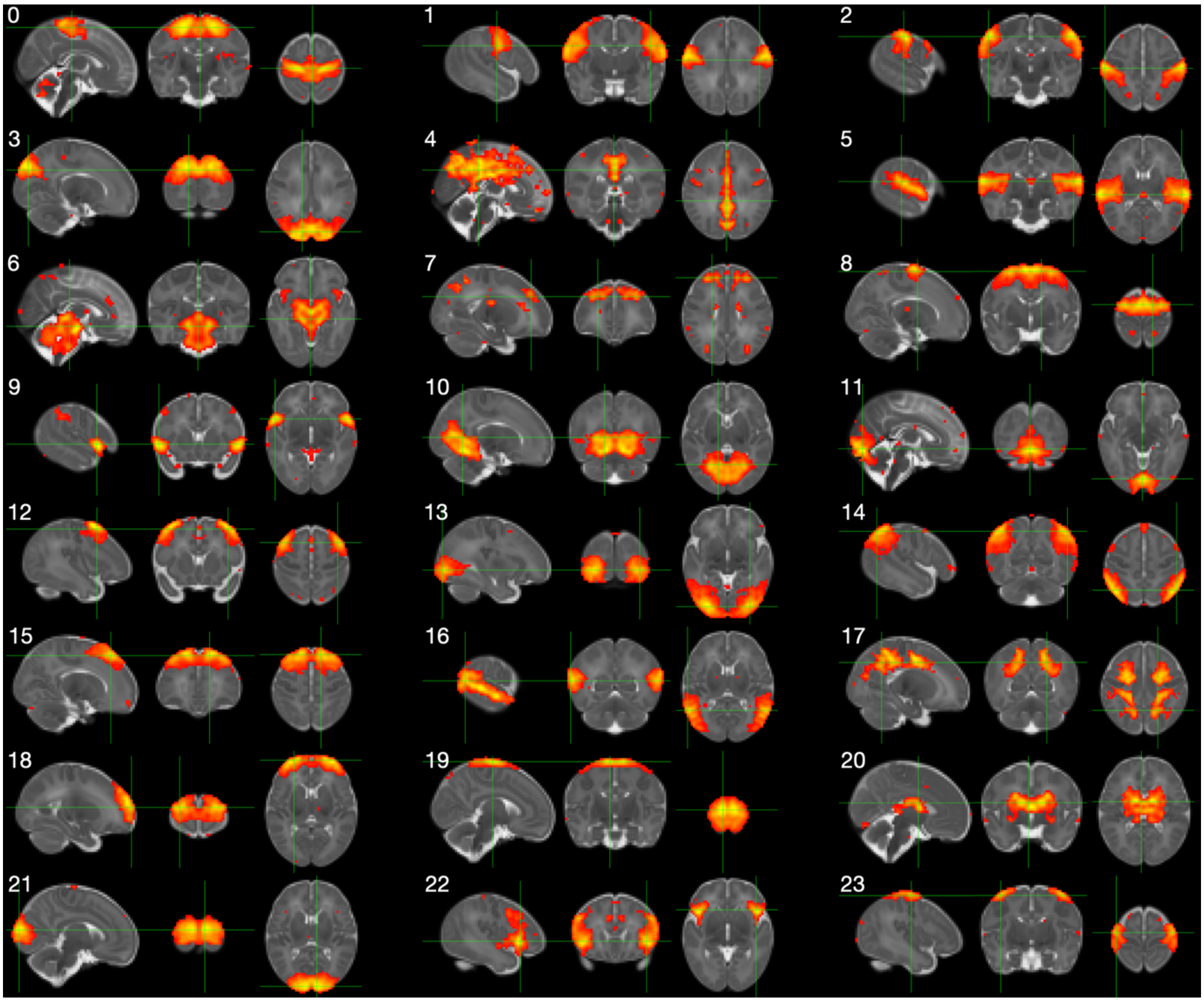

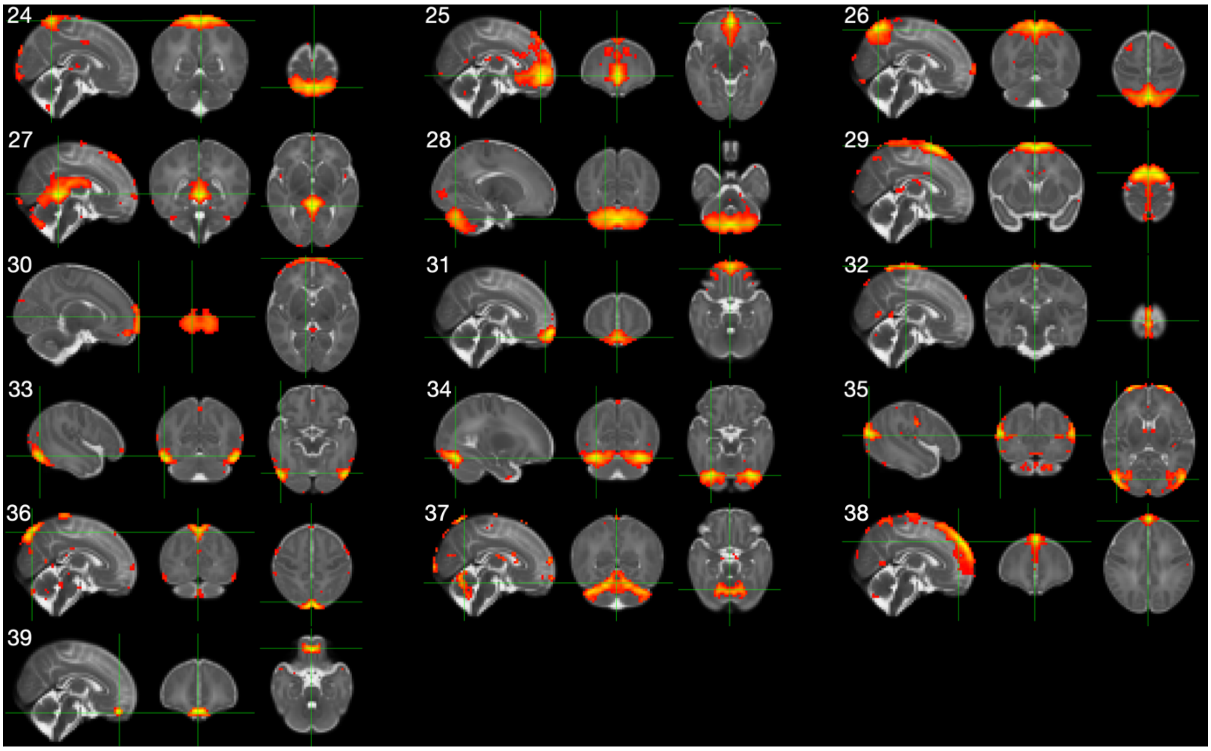

*Figure S 8. d = 40 neonatal group-ICA maps.*

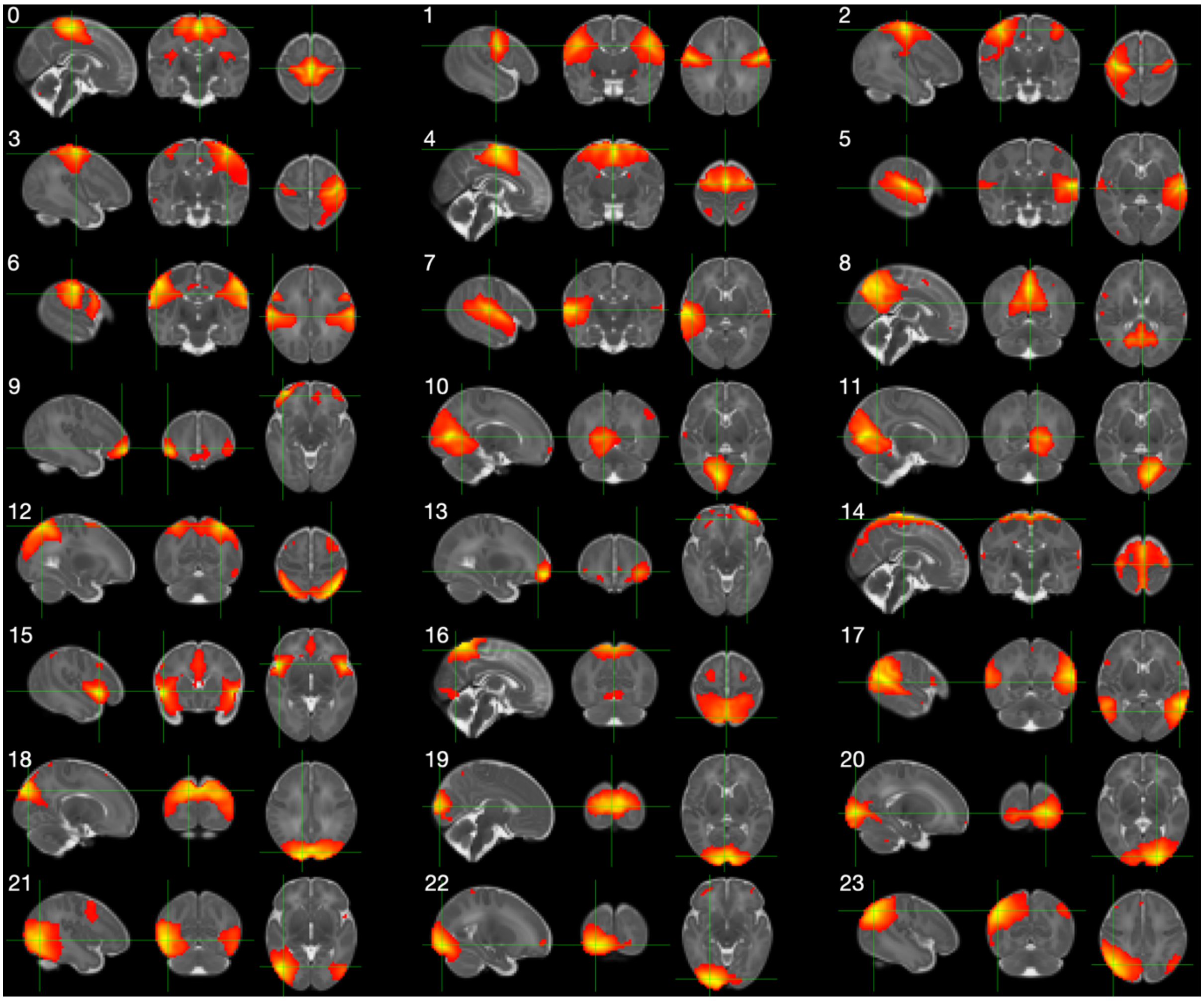

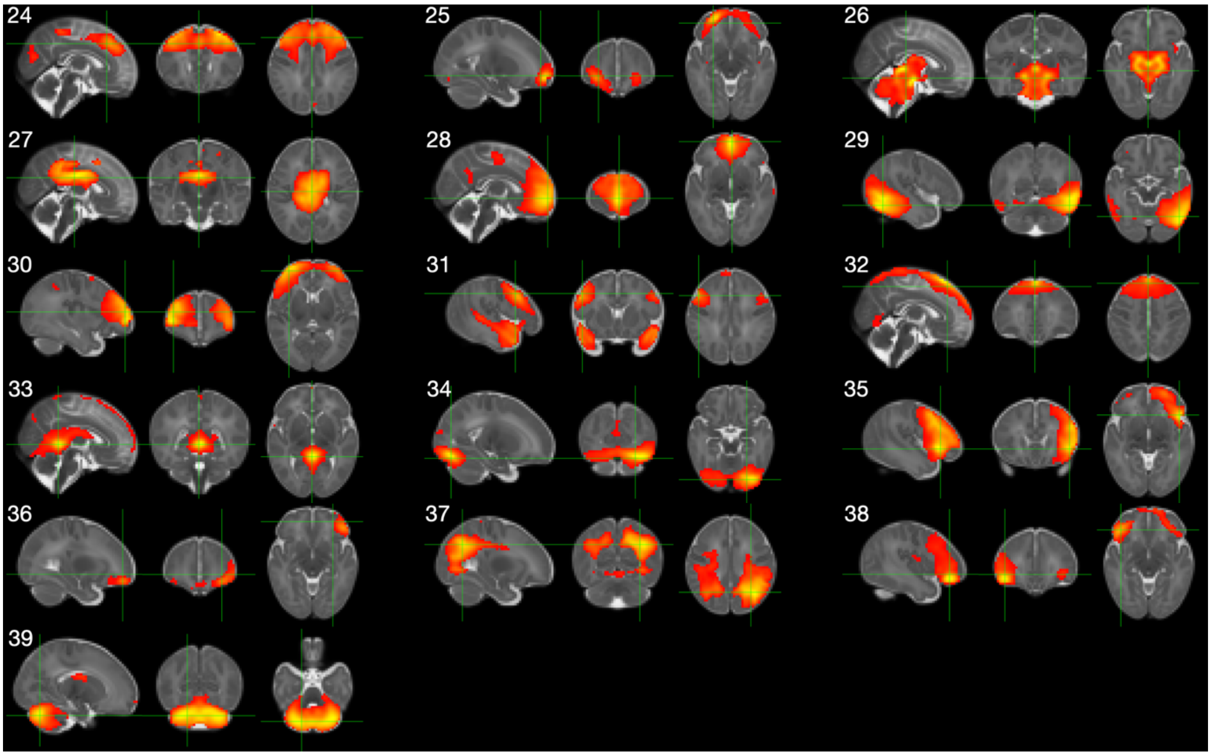

*Figure S 9. d = 40 fetal group-ICA.*

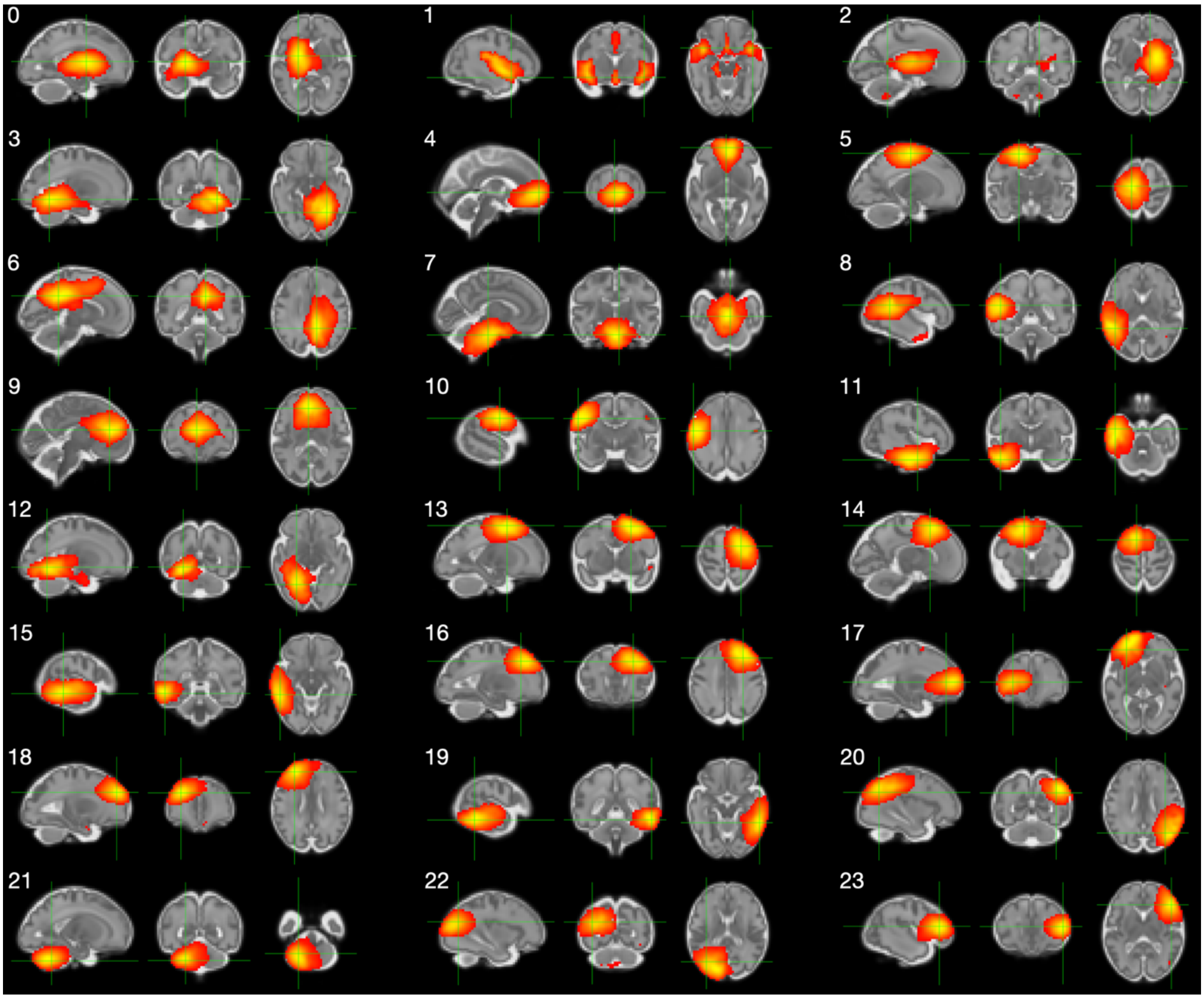

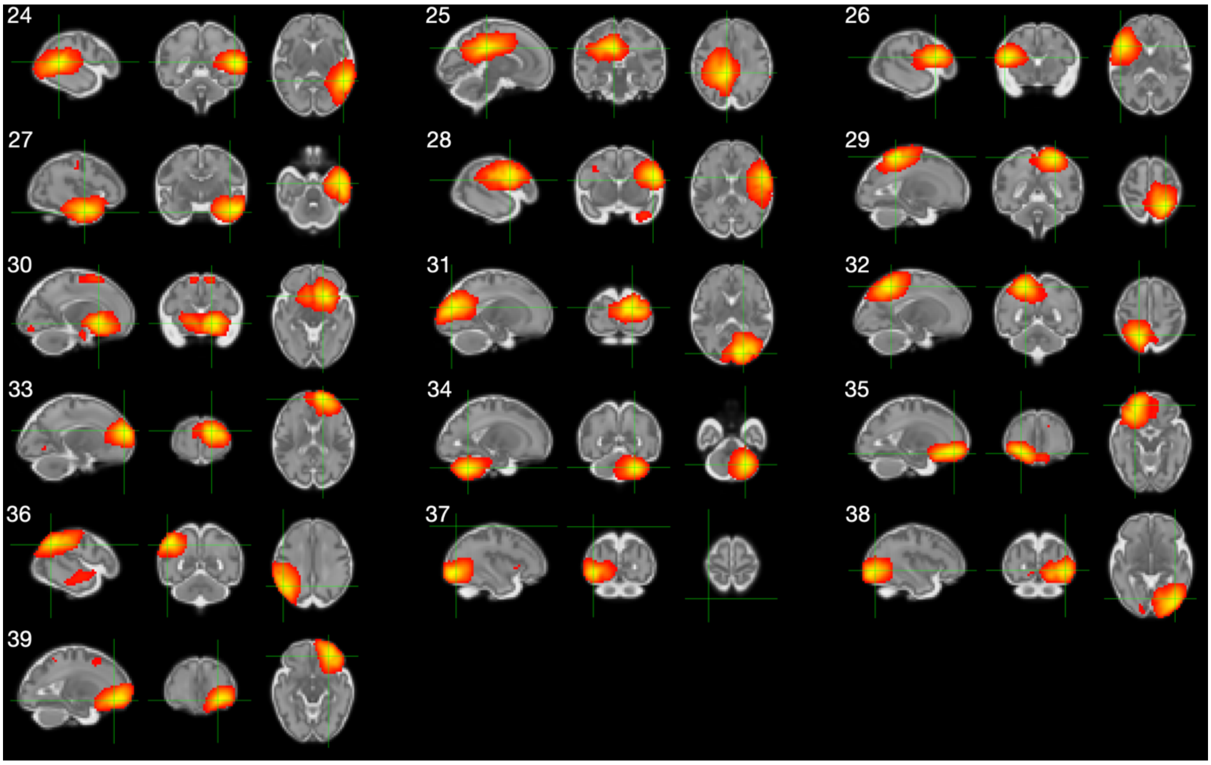

*Figure S 10. d = 40 left+right averaged thalamus-based fetal FCMs.*

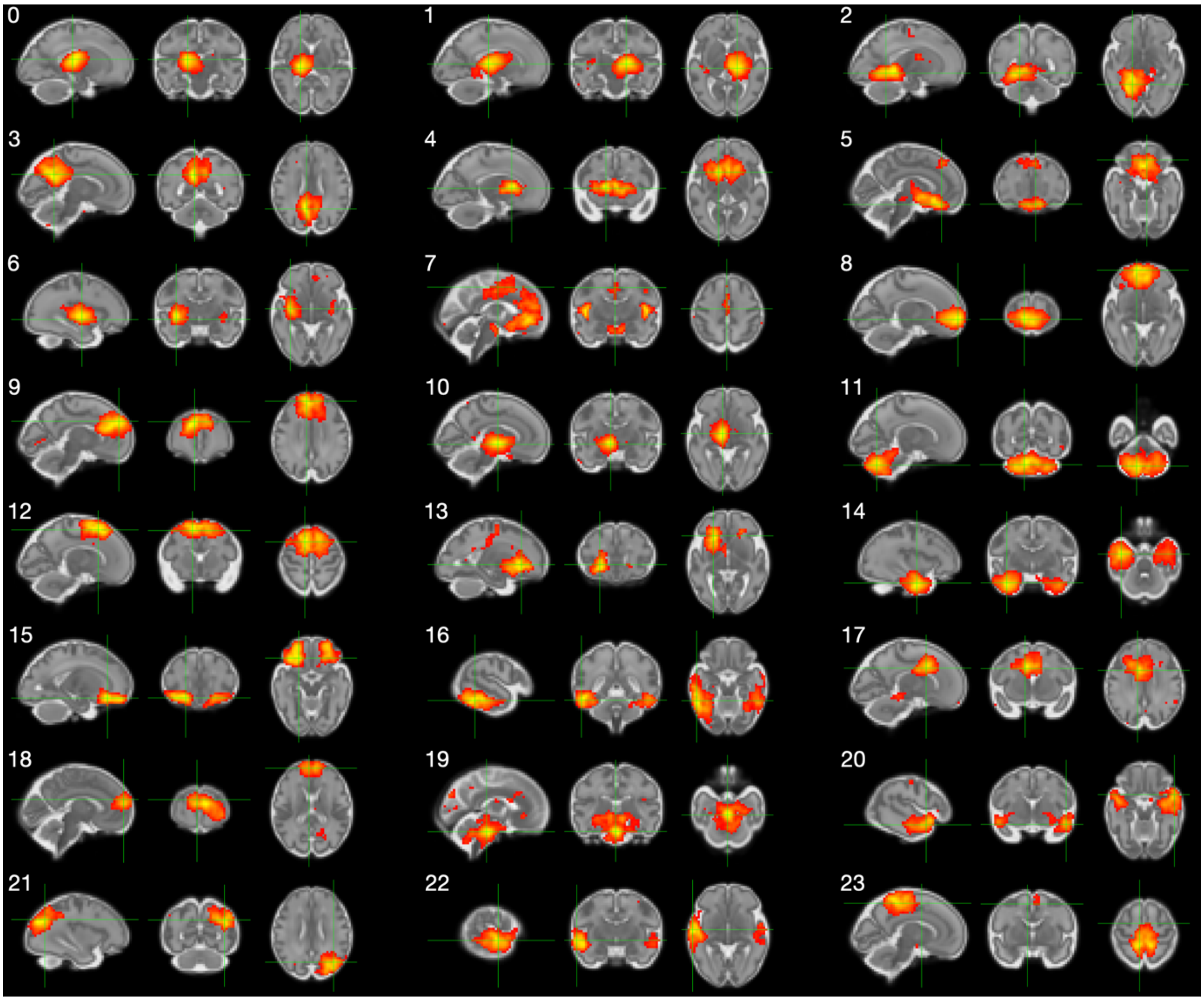

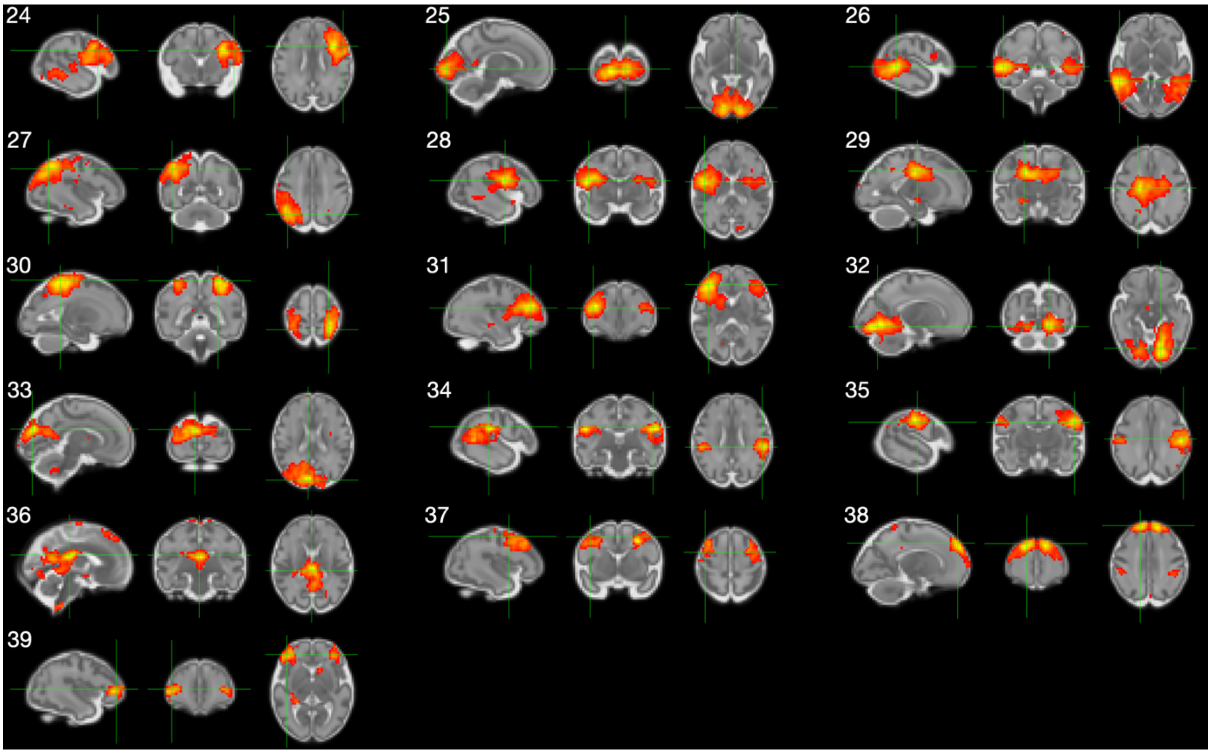

*Figure S 11. d = 40 fetal FCMs computed by concatenation of left thalamus and left cerebellum correlation maps.*

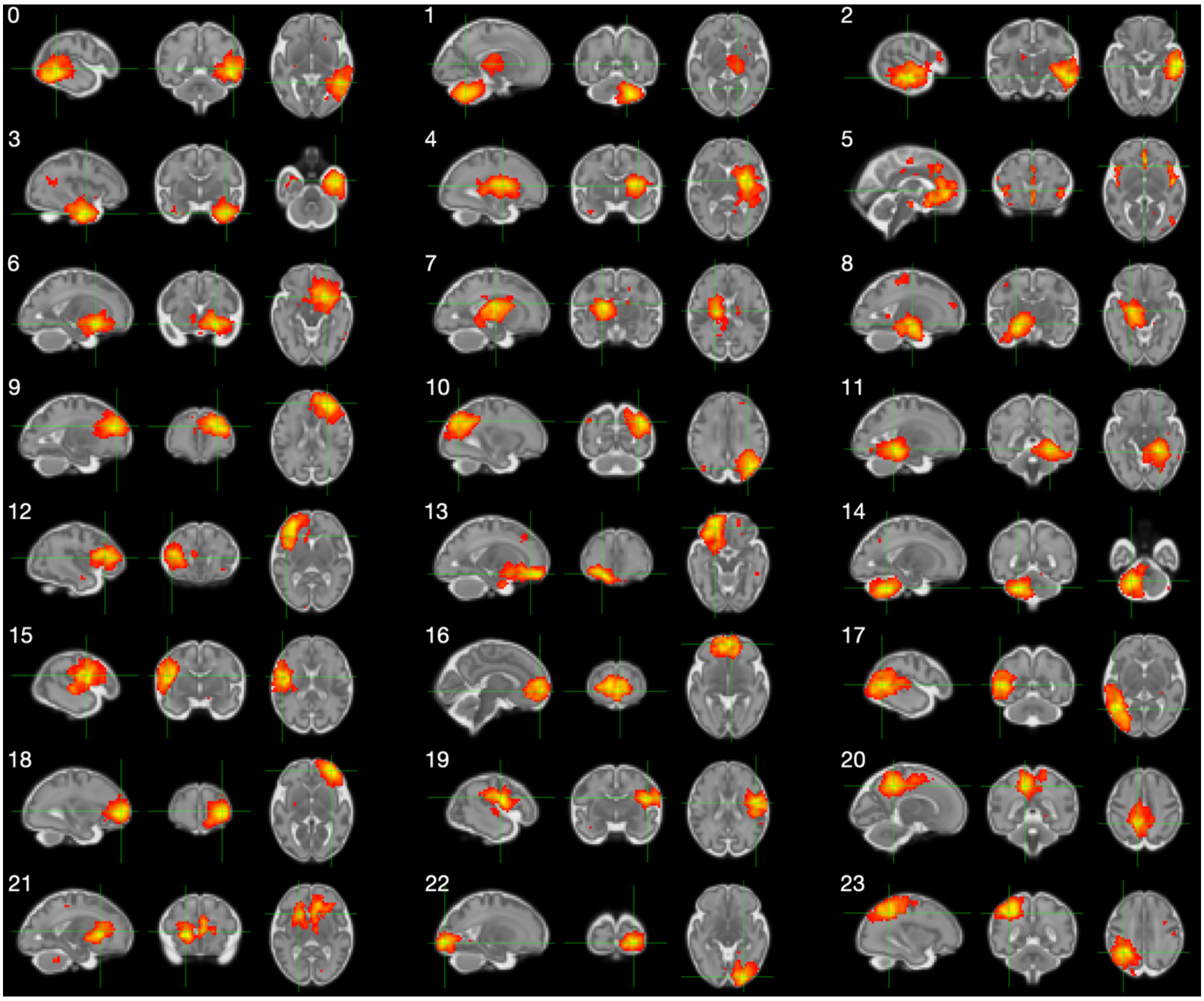

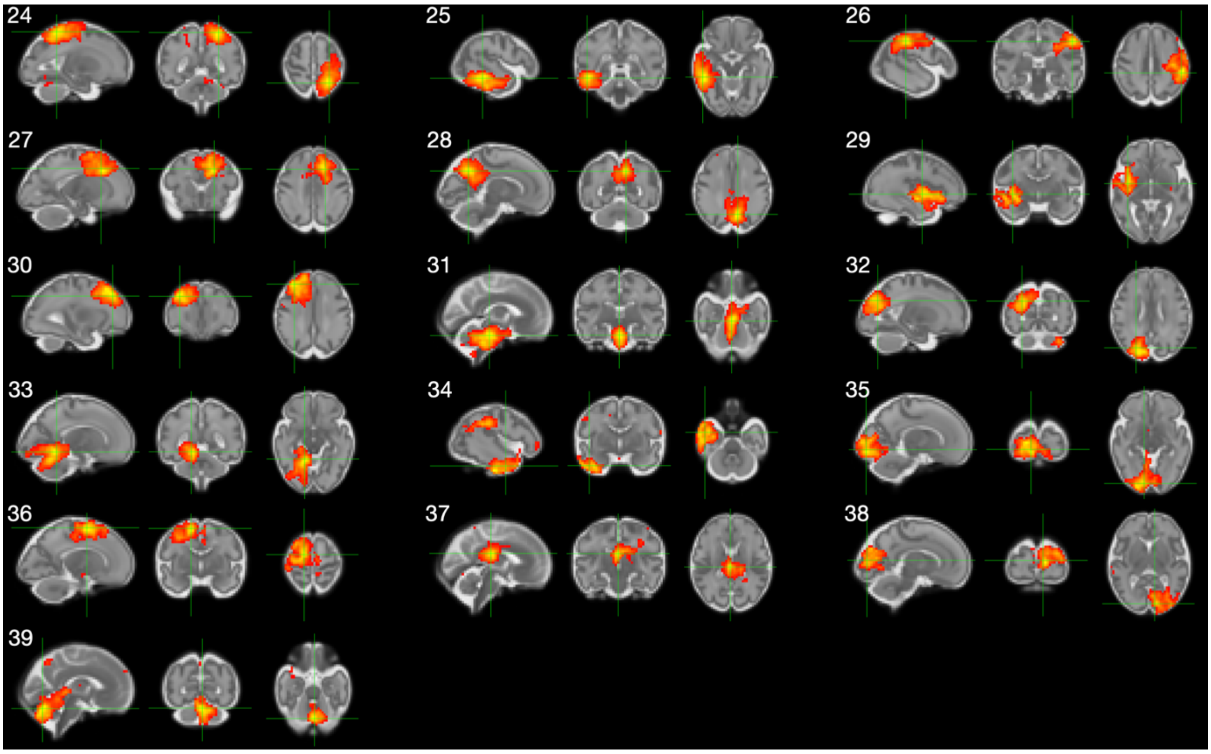

*Figure S 12. d = 25 fetal mrFCMs.*

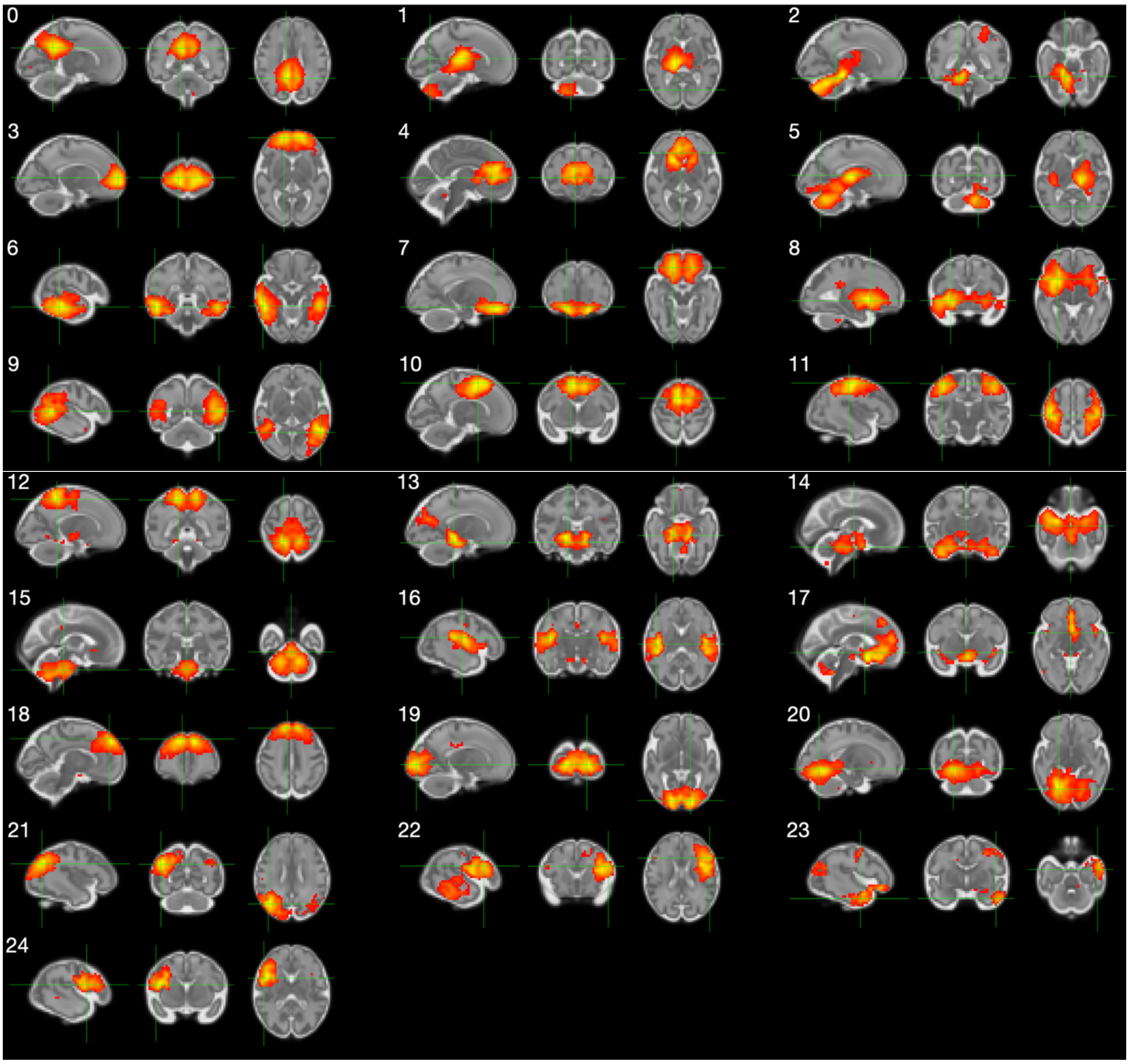
